# Calculating Muscular Driven Speed Estimates for *Tyrannosaurus*

**DOI:** 10.1101/2024.06.13.596099

**Authors:** Adrian T. Boeye, Scott Swann

## Abstract

Top speed estimates of extinct dinosaurs have been of long-standing interest to gain better understanding of the animals’ lifestyle and ecology. *Tyrannosaurus rex* top speeds have been examined using a wide range of methods that draw on more traditional biomechanical formulas, computer simulations, and allometric equations based on mass. However, these calculations may be made more precise using input from contemporary research on anatomy and biomechanics that account for mass allometry and scaling. This study builds on existing studies in anatomy, biomechanical data, and established equations for locomotion to calculate a muscular driven range of top speed for several (n=4) specimens that had sufficient data to undertake this work. When properly refined with additional data on muscle mass allometry and scaling, several adult specimens of *T. rex* could confidently be placed in a range of top speed from 7.7 to 10.5 m/s, and possibly up to 10.7 m/s. Additionally, a younger specimen of *T. rex* was analyzed and found to have a higher top speed than the adult *T. rex* at 6.3 to 14.5 m/s. Although the estimated top speeds in this study are slower than some previous estimates, these results find some support for slow running gaits and reinforce interpretations of *T. rex* as an active and effective apex predator. Future work can build upon this study by investigating how muscular driven top speeds may affect ontogenetic niche partitioning and prey species regularly targeted by adult *T. rex*.

## INTRODUCTION

A variety of methods have been used to estimate the maximum speed of extinct dinosaurs, from generalized formulas for vertebrates to more taxon-specific analyses (Alexander, 1976; Hutchinson and Garcia, 2002; Paul, 1998; Ruiz and Torices, 2013; Usami and Kinugasa, 2017; Thulborn 1990). Understanding the top speed of a predator holds a great deal of importance for the lifestyle and ecology as it can help to understand how active an animal is, as well as what animals it will regularly target (Carbone et al., 2011; Krauss and Robinson, 2013). Top speed estimates for *T. rex* have ranged from >13 m/s (Paul, 1998; Usami and Kinugasa, 2017) to a more conservative 5-11 m/s (Hirt et al., 2017; Hutchinson and Garcia, 2002; Sellers et al., 2017). The inability to examine the full range of physical characteristics in extinct animals often leaves key aspects of anatomy and biomechanics underexplored.

With such a wide range in estimated top speed (Fig. 1), quantitative analysis using more accurate anatomical and biomechanical data becomes essential to retrieve more precise results (Hutchinson, 2021). This study incorporates previously established methods (Alexander, 1976, Ruiz and Torices, 2013, Thulborn, 1990), relatively recent studies of anatomy (Bates et al., 2009, Hutchinson et al., 2005; Hutchinson et al., 2011), and biomechanics to calculate a plausible range of top speeds for *T. rex*. The methods of using precise inputs (Gatesy et al., 2009) into established equations results in a constrained and thus plausible range of muscular driven speed estimates.

**FIGURE 1.**
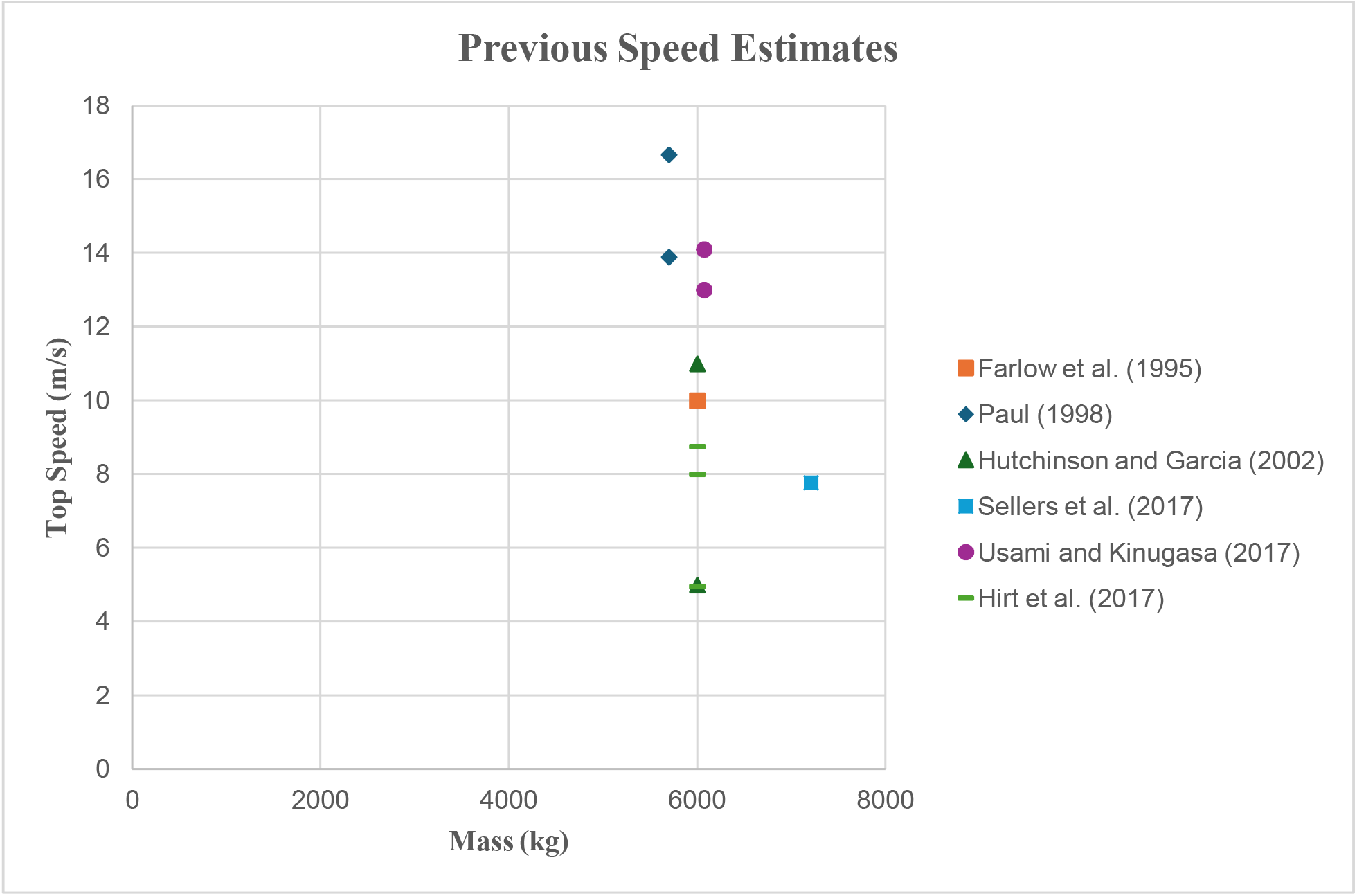
previous speed estimates, primarily based on MOR 555. Note that Paul (1998) used CMNH 9380 and Sellers et al. (2017) used exBHI 3033.

**FIGURE 2.**
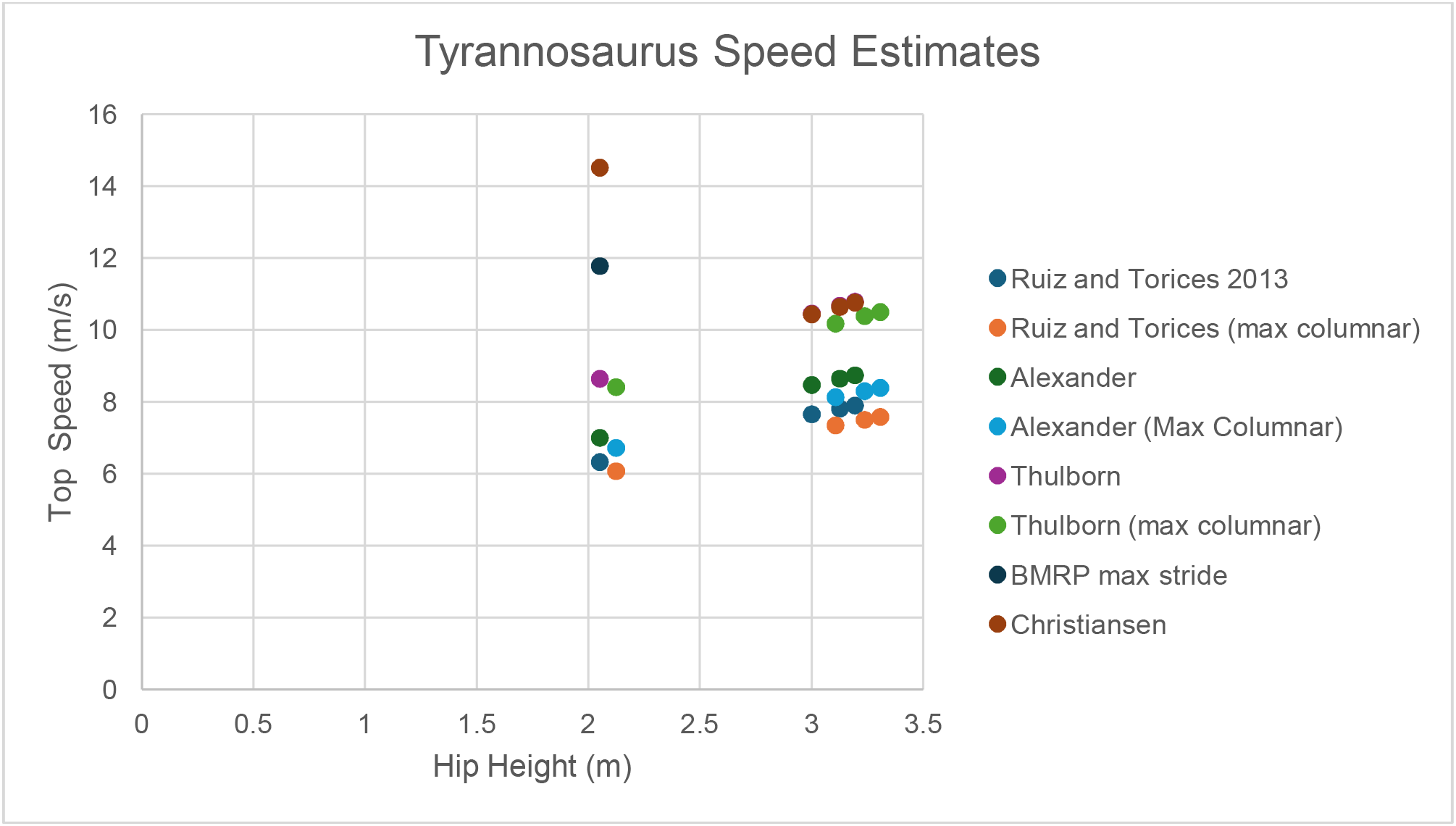
calculated speed estimates for Tyrannosaurus specimens BMRP 2002.4.1, MOR 555, exBHI 3033, and FMNH PR 2081.

While several equations exist that can calculate the speeds of animals based on hip height and stride length as derived determinants of the type of locomotion, which in turn would help to incorporate stride frequency as well as overall size (Alexander, 1976; Ruiz and Torices, 2013; Thulborn, 1990), without constrained and precise inputs the results these equations can produce are often implausible (e.g., Dececchi et al., 2020). This study uses data on midstance (Gatesy et al., 2009) and muscle mass (Bates et al., 2009; Hutchinson et al., 2011; Persons and Currie, 2011 Snively et al., 2019) to more precisely establish constraints and inputs for previously established formulas to produce a range of top speed. While such methods are not as complex as some recent computer simulations (Sellers and Manning, 2007; Sellers et al., 2009; Sellers et al., 2017), the approach and results from the present study are useful to help narrow down plausible speed estimates while remaining in line with current evidence. Another important consideration for speed that has been studied is the change in lifestyle undergone by adult *T. rex*, where the animal became larger and more robust as the animal aged (Carr, 2020). By analyzing several different *T. rex* specimens at different stages of ontogeny and testing them with several formulas to calculate top speed, this study also explores changes in speed and thus implied changes in niche across an animal’s life cycle.

## MATERIALS AND METHODS

### Calculations for top speed

Over many decades numerous methods have been developed and refined to investigate top speeds, including computer simulations (Sellers et al., 2007; Sellers et al., 2009; Sellers et al., 2017) and a variety of mathematical formulas (Alexander, 1976; Ruiz and Torices, 2013; Thulborn, 1990). Throughout all these works a universal importance is placed upon anatomy, morphology, and the effects of size (Biewener, 1989; Dick and Clemente, 2017; Hirt et al., 2017; Hutchinson and Garcia, 2002). This study uses several established mathematical formulas in conjunction with input guided by computer modelling (Gatesy et al., 2009) as well as insight from well-known mass related scaling principles (Dick and Clemente, 2017; Hirt et al., 2017).

For each specimen selected, four equations were utilized to calculate speed due to their ability to be adapted to account for changes in posture and by extension mass. Three of these equations are emphasized due to their ability to be adapted to better conform to scaling principles and can be found in Alexander (1976) with the formula v(m/s) = 0. 25(*g*)^0.5^× (SL)^1.67^× (□)^−1.17^ (Eq. 1), Ruiz and Torices (2013) where v(m/s) = 0. 226(g)^0.5^× (SL)^1.67^× (□)^−1.17^ (Eq. 2), and Thulborn (1990) where v(m/s) = [g□(SL/1. 8□)^2.56^]^0.5^ (Eq. 3). A fourth equation was also used following methods found in Christiansen (1998) that can be found in Supplementary file 1 (Eq. 4), however the results should be taken with caution for animals above 6 tonnes since the original study does not include any animals above said size range. V is velocity in m/s, SL is stride length in meters, and *h* being hip height in meters. It should be noted that hip height will not be directly equivalent to the actual leg length of the animal, primarily due to changes in posture as animals move (Dick and Clemente, 2017; Gatesy et al., 2009). When calculating stride length, the length of the leg to the third metatarsal was measured, due to this part of the leg being the first point of contact between a moving animal and the ground (Rothschild, 2001), before being multiplied by a constant determining the type of locomotion found in Thulborn (1990). For animals engaged in a run this constant would need to be greater than 2.9 (minimum of 2.91), resulting in SL being leg length to the third metatarsal × 2.91. For animals engaged in a walk this constant would be 2.9 or less (SL= leg length to the third metatarsal × 2.9).

### Muscle mass allometry

Additional analysis was conducted on the muscle mass allometry of several different specimens of *T. rex* (see supplementary file 1). These calculations were broken into 3 sections: total leg muscle mass vs body mass, total leg muscle mass and included m. Caudofemoralis (mCF) muscle mass vs body mass, and total leg muscle mass and an included 40% increased mCF muscle mass following Snively et al. (2019) vs body mass. When using the enlarged mass derived from Snively et al. (2019), the additional muscle mass was also added to the original body mass (see supplementary file 1). While other studies such as Bishop et al. (2021) do not include the mCF in allometry models of sauropsids, other work strongly suggests that this muscle was key to the locomotion of these animals (Persons and Currie, 2011; Snively et al., 2019). As such, including this additional muscle mass may provide additional insight. Ultimately both methods are useful and may help with future comparisons and are included in Supplementary file 1.

### Anatomical data

Anatomical data was obtained from several sources, including Bates et al. (2009), Hartman (2013), Hutchinson et al. (2011), Persons and Currie (2011), Persons and Currie (2016), and Snively et al. (2019). When studying muscle mass allometry Bates et al. (2009) provides detail on total muscle mass in the leg but does not include information on the size of the mCF and as such a reconstructed value derived from Persons and Currie (2011) is used, maintaining the same ratio of mCF mass to total body mass outlined in the work (a 348 kg mCF compared to a 6000 kg *T. rex*). One point of interest is the exact ratio of body mass to mass of the mCF in Persons and Currie (2011). The original paper uses a 4.5 tonne estimate for exBHI 3033, a likely underestimate (Hutchinson et al., 2011). A possible alternative interpretation found in Persons and Currie (2011) is to use their muscle mass restoration of 261 kg with a body mass of 7.6 tonnes derived from Bates et al. (2009). If this is done, then the overall ratio of muscle mass to body mass drops to levels like Hutchinson et al. (2011). For information regarding these calculations see Supplementary file 1. A brief note should be made for Persons and Currie (2011) that the estimate includes both the mCaudofemoralis Longus and mCaudofemoralis Brevis, resulting in a slightly higher estimate than Hutchinson et al. (2011) which exclusively measures the mCaudofemoralis Longus. When using muscle mass allometry in Hutchinson et al. (2011) data is provided for the total amount of extensor muscle in the legs and the mCFL. Since extensor muscle composes 70% of total muscle mass in the leg (Paul, 2008), the value calculated in total muscle mass allometry is adjusted to include the total muscle mass.

Data for the legs of the various animals was collected from several sources, including Hartman (2013), Hutchinson et al. (2011), and Persons and Currie (2016). For determining the midstance of the animal, data from Gatesy et al. (2009) was used, with the total leg length of all specimens being based on the ratio of MOR 555 found in Gatesy et al. (2009). The midstance itself is based again on Gatesy et al. (2009) with a normal midstance being 3m for MOR 555 and a similar ratio being maintained for other *T. rex* specimens, while the maximum hip height being 95% of the total leg length. This height should be taken with some caution however as it would likely be quite uncomfortable for the animal and could only be achieved with support imparted by soft tissue beneath the foot.

### Specimens

The following specimens were selected for analysis in this work, including MOR 555, exBHI 3033, FMNH PR 2081, and BMRP 2002.4.1. These specimens were selected due to the completeness of their fossils as well as the significant amount of research focused on these specimens (Bates et al., 2009; Hutchinson et al., 2011). Some additional attention must be given to exBHI 3033, FMNH PR 2081 and BMRP 2002.4.1, primarily due to available data and reference material. For exBHI 3033 there is significant variation in the total mass estimates between Bates et al. (2009) and Hutchinson et al. (2011), approximately 6 and 7.6 tonnes respectively. FMNH PR 2081 is restored as weighing 9.5 tonnes in Hutchinson et al. (2011), however a notably lighter restoration at 8.4 tonnes is also outlined in Snively et al. (2019). Data is given for the muscle mass of the leg (Snively et al., 2019) and since the primary discrepancy between the leaner and heavier models of Snively et al. (2019) is primarily in the ribcage, use of said leg muscle mass is appropriate. When restoring the muscle of the tail a ratio like the one found in Persons and Currie (2011) was used. When examining BMRP.2002.4.1, the speed estimates must be taken with caution owing to the lack of data available to reconstruct a proper gait for immature *T. rex*.

### Considerations for assessing speed estimates

When assessing which speed estimates are most likely, a few points must be considered. First is center of mass (COM), which as Gatesy et al. (2009) describe is key to a *T. rex*’s ability to run, with a center of mass positioned more than 0.5m anterior to the hip resulting in an inability to run. COM estimates can be found in Bates et al. (2009), Hutchinson et al. (2011), and Snively et al. (2019). Second, overall body mass: according to scaling principles heavier animals are typically going to be slower than lighter ones (Dick and Clemente, 2017; Hirt et al., 2017). Third is muscle vs body mass allometry; animals which demonstrate greater allometry are likely to be faster than animals with a lower muscle mass allometry. However, a caveat should be made when examining allometry and body mass; if an animal is significantly lighter than another animal it is likely to be faster than its heavier even if the heavier animal demonstrates somewhat more positive allometry. For a heavier animal to be faster than its lighter counterpart, it would need to demonstrate significantly greater positive muscle mass allometry. Fourth, the various needs to support mass will shift and become greater as an animal gets heavier. As such it is more likely that heavier animals will either need to shift their stance to become more columnar or simply become slower, both consequences of their greater mass (Dick and Clemente, 2017).

INSTITUTIONAL ABBREVIATIONS — **MOR**, Museum of the Rockies, Bozeman, Montana U. S. A; **BHI**, Black Hills Institute, Hill City, South Dakota, U. S. A; **FMNH**, Field Museum of Natural History, Chicago, Illinois, U. S. A; **BMRP**, Burpee Museum of Natural History, Rockford, Illinois, U. S. A; **CMNH**, Carnegie Museum of Natural History, Pittsburgh, Philadelphia, U.S. A.

ANATOMICAL ABBREVIATIONS— **mCF**, musculus Caudofemoralis; **mCFB**, musculus Caudofemoralis Brevis; **mCFL**, musculus Caudofemoralis Longus

## RESULTS

To see the complete range of estimates for muscle mass allometry and speed for each specimen using each type of gait, see Supplementary file 1. The lightest specimen, BMRP 2002.4.1 displayed a wide range in top speed estimates from 6.3 to 14.5 m/s as calculated by the selected equations. Muscle mass allometry ranged from 0.811 to 0.868. With a COM anterior to the hip by 0.276m (Hutchinson et al. 2011), gait reconstruction done by Gatesy et al. (2009) suggests that BMRP 2002.4.1 could enter a true run. If a larger mCF was present (Snively et al. 2019) this COM would be shifted further posteriorly. A true run would tighten the range of speed to 8.7 to 14.5 m/s.

The second lightest specimen, MOR 555 displayed a range of speed between 7.7 and 10.5 m/s. Muscle mass allometry ranged from 0.856 to 0.912. This allometric range is, in part, due to the differences in muscle mass of the leg between Bates et al. (2009) and Hutchinson et al. (2011) as well as the differences in the estimated mCF derived from Persons and Currie (2011) and Hutchinson et al. (2011). COM also demonstrated variability across reference material; Bates et al. (2009) restored a COM 0.468m anterior to the hip while Hutchinson et al. (2011) restored a COM 0.572m. If the Bates et al. (2009) restoration is used then MOR 555 could enter a true run, while the restoration from Hutchinson et al. (2011) could not. This could change, however, with the enlarged mCF, since Bates et al. (2009) performed tests with several animals in the study that enlarged the caudal sections of the tested animals. When the caudal section of MOR 555 was increased 7.5% and anterior elements reduced 7.5% the COM shifted to 0.295m. If the mCF is enlarged as suggested by Snively et al. (2019), then the COM may shift posteriorly of the 0.5m anterior point from the hip in turn allowing the restoration found in Hutchinson et al. (2011) to enter a true run.

The third tested specimen, exBHI 3033 displayed a range of speed between 7.8 and 10.7 m/s. Muscle mass allometry ranged from 0.859 to 0.906. Like MOR 555, COM ranged between Bates et al. (2009) and Hutchinson et al. (2009). The Bates et al. (2009) model possessed a COM of 0.572m anterior of the pelvis, with their enlarged tail model shifting this COM to 0.385m anterior to the hip and may enable running. It should be mentioned, however, that this model of exBHI 3033 approximately 1.5 tonnes heavier than the Hutchinson et al. (2011) model and as such other principles of scaling may need to be accounted for with the increased mass which could result in running being impossible (Dick and Clemente, 2017; Hirt et al., 2017; Hutchinson, 2021). The Hutchinson et al. (2011) model possessed a COM 0.524m anterior to the hip. An enlarged mCF would likely shift the COM posteriorly of the 0.5m anterior point and thus enable running.

The fourth and heaviest specimen, FMNH PR 2081 displayed a range of speed between 7.9 and 10.8 m/s. Muscle mass allometry ranged between 0.807 and 0.875. The lean model from Snively et al. (2019) did not have any listed COM, and as such running ability is unknown but improbable due to its mass and larger anterior section of the body (Hutchinson et al. 2011; Hutchinson, 2021). The Hutchinson et al. (2011) model was restored with a COM 0.801m anterior to the hip, eliminating any ability to run. Even with an enlarged mCF, FMNH PR 2081 was unlikely to have the COM shifted posteriorly beyond the 0.5m anterior to the hip point.

When accounting for the improbability of running, the range of speed for FMNH PR 2081 is between 7.9 and 8.7 m/s.

## DISCUSSION

All *T. rex* specimens displayed a fair range of speed, however most of the adult specimens tended to cluster around an intermediate estimate close to other results produced by Sellers et al., (2007). Some of the higher end estimates for the adult *T. rex* do fall above the most recent studies that quantified the top speed of *T. rex* (Hirt et al., 2017, Sellers et al., 2017). This is largely due to different approaches and methods of calculation, with the current study focusing on a muscle driven model with strict inputs on anatomy. The other studies with lower top speed estimates are invaluable in providing insight into potential limitations on speed, and like this study, reaffirm previous work that suggests *T. rex* was unlikely to run faster than 11 m/s (Hutchinson and Garcia, 2002). Ultimately, when studying an animal like *T. rex* there will be a great deal that will never be known, but together these works, this study included, help to remove some of these unknowns (Hutchinson, 2021).

When interpreting the reliability of the results, some nuance must be used. Eq 1 to 3 are all relatively reliable due to their ability to be fine-tuned for the animal’s midstance, but it should still be noted that these equations can overestimate the speed of large animals (e.g., Dececchi et al., 2020). Eq. 4. should be taken with the most caution, however, since it cannot be scaled for animals beyond 6 tonnes, and indeed higher estimates are seen due to a lack of this scaling (see supplementary file 1). Additionally, when looking at midstance additional attention should be dedicated to the potential for the maximum columnar stance to better support larger animals (Dick and Clemente, 2017). While Gatesy et al. (2009) find to be a preferred midstance for MOR 555, heavier animals may need to resort to a more columnar stance to gain more support (Biewener, 1989; Dick and Clemente, 2017) With these caveats in mind, these estimates have all been recovered before in previous work (Hutchinson and Garcia, 2002) and as such are likely to be plausible. A particular point of interest is MOR 555, which has had an exceptional amount of work performed on it. Given that both Eq. 3 and Eq. 4 recover nearly identical ranges of speed (10.5 m/s) when using precisely guided inputs (Bates et al., 2009; Gatesy et al., 2009; Hutchinson et al., 2011) it is quite likely that if the most recent restorations are accurate (Snively et al., 2019) and MOR 555 could run, then a top speed of 10.5 m/s is likely.

Additional time should be taken to analyze exBHI 3033 due to its differing size and stage of ontogeny (Carr, 2020). In the case of exBHI 3033, there is a large deal of variation in size from 6 tonnes (Hutchinson et al. 2011) to 7.5 tonnes (Bates et al., 2009). As discussed in the results the COM is anterior of the 0.5m limit determined by Gatesy et al. (2009) and would eliminate exBHI 3033’s ability to run. However, if the mCF is enlarged as suggested by Snively et al. (2019) the COM would be posteriorly of the 0.5m point and allow the animal to enter a slow run. This possibility is especially interesting since Hutchinson et al. (2011) restore exBHI 3033 as more muscular than the slightly lighter MOR 555. Given the slightly greater muscle mass allometry and only marginally greater mass, it is very possible that Hutchinson et al. (2011) restoration of exBHI 3033 was the fastest of the adult *T. rex*. This is especially likely if the larger estimates of the mCF continue to be accepted. The heavier Bates et al. (2009) model does possess a COM posterior of the 0.5m anterior point. Additional analysis would need to be conducted to determine the constraints imparted by this heavier size, and until such work is done the running ability of a heavier exBHI 3033 remains ambiguous.

FMNH PR 2081 also is of interest, particularly due to its massive size. The “lean” estimate contains a great deal of unknowns, most prominently in the lack of an established COM. While the lean model is certainly lighter than the restoration in Hutchinson et al. (2011), it still shows relatively reduced muscle mass allometry compared to other specimens of *T. rex*. Given this information it is likely that the lean model of FMNH PR 2081 was likely incapable of running. A mass of 8.4 tonnes would be unable to generate the necessary force to enter a running gait. These assertions are also supported by the Hutchinson et al. (2011) model which has much less ambiguity. The Hutchinson et al. (2011) model is a full tonne heavier than the lean model and shows a COM well past the 0.5m point, as well as displaying the lowest muscle mass allometry of any of the sampled *T. rex*. Additionally, Hutchinson et al. (2011) outline some key differences between FMNH PR 2081 and the lighter *T. rex* in their study, including a proportionately larger torso, proportionally smaller legs and tail, as well as a larger head. Given these data it is reasonably confident to assert that FMNH PR 2081 could not run and was entirely restricted to fast walking.

The results of this work provide some key insight into a variety of different aspects of speed, and ultimately lifestyle throughout ontogeny. Starting with muscle mass allometry, it is clear that while *T. rex* were exceptionally muscular (Hutchinson et al. 2011), all values of allometry retrieved were moderately negative, especially when compared to smaller contemporary animals (Bishop et al. 2 021). This strongly suggests that adult *T. rex* were slower than lighter animals and bolsters the general cluster towards a slow to intermediate speed (Dick and Clemente, 2017; Hirt et al., 2017). However, this somewhat lower allometry does not affect the smaller and lighter BMRP 2002.4.1 as significantly. Contrary to this point, the use of Eq. 3 and Eq. 4 in this work has recovered BMRP 2002.4.1 as potentially faster than any adult *T. rex*. The lower muscle mass allometry suggests that while the animal may have been slower than similar sized cursorial animals of today, it was still a relatively quick animal.

Another interesting result of this study is in the general shift in locomotor performance. Slower speeds have been suggested for older and heavier *T. rex* adults (Hirt et al., 2017; Hutchinson et al., 2011), and when precise constraints are used, this work also supports these theories. Recently, proposed changes in lifestyle have been suggested for *T. rex* as the animal aged with younger and faster animals targeting different prey (Carr, 2020; Rowe and Snively, 2022; Therrien et al., 2023). The changes in speed found by this work do provide some support to this, while acceleration would also likely be a key component to hunting success, it seems likely that younger *T. rexes* were better able to keep pace with the smaller animals of their environment.

While slower than their younger counterparts, adult *T. rex* could still pursue a diverse range of prey. Current suggestions of ontogenetic niche partitioning propose that adult *T. rex* hunted the larger animals of their environment (Holtz, 2021; Rowe and Snively, 2022). While the largest animals such as *Edmontosaurus, Triceratops*, or *Ankylosaurus* were likely pretty close to an adult *T. rex* in speed (Hirt et al., 2017; Paul and Christiansen, 2000), and in some cases likely being slower than a *T. rex*, it is highly unlikely that *T. rex* targeted the largest animals it could. While *T. rex* could bring down these animals, such attacks would pose an unnecessarily high risk to the attacking *T. rex*. Instead, it would likely target younger, sick, old, or weak animals. In the case of these younger animals’ speed and acceleration would be vital, especially since some of these younger animals were quite fast (Sellers et al., 2009). While this work does not calculate acceleration, it does support the possibility that an adult *T. rex* could briefly keep pace with some of these animals before catching up to and subduing them with a powerful bite. If restorations with a COM posterior of the 0.5m point are favored true running remains viable then an adult *T. rex* would better be able to keep up with these smaller and lighter animals. Tentatively this work finds potential support for a true running model which would align with selection pressure placed on adult *T. rex* and their need to catch prey. Future work could build upon these findings by examining how selection pressure and prey capture would force certain locomotor demands on *T. rex* to capture prey. In this regard, studies in the ability of *T. rex* to accelerate would be extremely useful.

## CONCLUSION

While the true speed of *T. rex* will never be certain, the results from this study lend support to interpretations of this theropod as a reasonably fast and active predator. The top speed ranges found in this study help to narrow down this range of estimates and fall within the range of previous quantitative studies that focused on musculature. This study used well-established equations for locomotion in conjunction with precise biomechanical and anatomical inputs (Bates et al., 2009, Gatesy et al., 2009; Hutchinson et al., 2011; Persons and Currie, 2011; Snively et al., 2019). The findings of this work support a reasonably quick adult *T. rex* and favor the potential for lighter adult *T. rex* to engage in slow running less than 11 m/s. Additionally, this study offers another line of evidence for ontogenetic niche partitioning during the life cycle of *T. rex*. Though not as fast as other interpretations (e.g., Paul, 1998; Usami and Kinugasa, 2017), the intermediate values obtained in this study are still reasonably fast for such a massive animal. Proposed speeds here would certainly be fast enough to capture prey by a variety of means, supporting interpretations of *T. rex* as an active apex predator (Carbone et al., 2011; Krauss and Robinson, 2013).

## SUPPLEMENTARY FILES

Supplementary_File_1.xlsx: Raw data including body mass estimates, leg and mCF muscle mass estimates, muscle mass allometry, data on limbs, data on posture, and calculations of speed.

## Notes

### Competing Interest Statement

The authors have declared no competing interest.

